# *SUFI:* An automated approach to spectral unmixing of fluorescent multiplex images captured in mouse and postmortem human brain tissues

**DOI:** 10.1101/2021.01.28.428639

**Authors:** Vijay Sadashivaiah, Madhavi Tippani, Stephanie C. Page, Sang Ho. Kwon, Svitlana V. Bach, Rahul A. Bharadwaj, Thomas M. Hyde, Joel E. Kleinman, Andrew E. Jaffe, Kristen R. Maynard

## Abstract

Multispectral fluorescence imaging coupled with linear unmixing is a form of image data collection and analysis that uses multiple fluorescent dyes - each measuring a specific biological signal - that are simultaneously measured and subsequently “unmixed” to provide a read-out for each individual signal. This strategy allows for measuring multiple signals in a single data capture session - for example, multiple proteins or RNAs in tissue slices or cultured cells, but can often result in mixed signals and bleed-through problems across dyes. Existing spectral unmixing algorithms are not optimized for challenging biological specimens such as postmortem human brain tissue, and often require manual intervention to extract spectral signatures. We therefore developed an intuitive, automated, and flexible package called *SUFI*: spectral unmixing of fluorescent images (https://github.com/LieberInstitute/SUFI). This package unmixes multispectral fluorescence images by automating the extraction of spectral signatures using Vertex Component Analysis, and then performs one of three unmixing algorithms derived from remote sensing. We demonstrate these remote sensing algorithms’ performance on four unique biological datasets and compare the results to unmixing results obtained using ZEN Black software (Zeiss). We lastly integrate our unmixing pipeline into the computational tool *dotdotdot* that is used to quantify individual RNA transcripts at single cell resolution in intact tissues and perform differential expression analysis of smFISH data, and thereby provide a one-stop solution for multispectral fluorescence image analysis and quantification. In summary, we provide a robust, automated pipeline to assist biologists with improved spectral unmixing of multispectral fluorescence images.

## Introduction

Multispectral fluorescence imaging and linear unmixing is a powerful approach for visualizing and quantifying multiple molecular properties of tissues and cells in a single experiment. In fluorescence microscopy, the intensity value at each pixel is proportional to the photoemission of fluorophores (1). Spectral imaging extends this approach by recording pixel intensity values at multiple wavelength bands across the electromagnetic spectrum (2). For each pixel, spectral unmixing (SU) aims to recover the material source (endmembers) and the proportion of each material (abundances). Since the fluorescent light emissions mix linearly (3), individual signals can be mathematically disentangled based on the relative contribution of each spectral signature (also known as a reference emission profile or “fingerprint” or “endmember”) present in the image (3,4) in a process called “linear unmixing.” Linear unmixing can distinguish fluorophores with similar emission spectra (2,5) and effectively remove background noise and autofluorescence from the fluorophore signal (3,6). One such autofluorescent material is lipofuscin, a yellow-brown pigment granule composed of lipid-containing residues of lysosomal digestion, which is highly expressed in postmortem human brain tissue and poses a major challenge for fluorescent imaging (7).

However, existing approaches for spectral unmixing, such as linear unmixing (4), similarity unmixing (8), LUMoS unmixing (9), are limited and have not been well optimized for assays that generate punctate signal, such as single molecule fluorescent in situ hybridization (smFISH,) or complex tissue specimens containing abundant lipofuscin autofluorescence, such as postmortem human brain. Linear unmixing can be performed using proprietary software that accompanies the microscope used for image acquisition, for example LSM780 microscope/Zen software (Zeiss) and Vectra Polaris Imaging System /Inform software (Akoya Biosciences), but this leads to several potential weaknesses as the exact algorithms used for unmixing are often proprietary, creating a potential “black box” in the data processing pipeline. Another major challenge of existing linear unmixing approaches is that users are often required to create individual reference spectrum before unmixing, which is laborious and may or may not be relevant for the particular image under study. Manual fingerprint generation is also prone to error and user bias since individual pixels need to be selected by the experimenter. Lastly, and perhaps most practically, linear unmixing of large brain sections captured in 4 dimensions (x, y, z, lambda) and requiring unmixing of 6 fluorescent channels (e.g. 4-plex smFISH, nuclear stain, lipofuscin autofluorescence) is computationally intensive. Many softwares also do not allow for batch processing and each image must be unmixed individually. This process is particularly cumbersome for large-scale datasets containing several hundred images that need to be unmixed.

Multiplex single-molecule fluorescence in situ hybridization (smFISH) using RNAscope technology (Advanced Cell Diagnostics) has emerged as a powerful approach for localizing and quantifying gene expression at cellular resolution in mouse and human brain tissues (10,11). Complementary to other next generation sequencing technologies, smFISH allows measurement of cell-to-cell variability in gene expression (12,13) and intracellular localization of mRNA transcripts (14). The widespread generation of single-cell RNA-sequencing (scRNA-seq) data sets in the brain has fueled a resurgence of multiplex smFISH to validate cell-type-specific molecular profiles by visualizing individual transcripts at cellular resolution in brain sections (15,16). Combinatorial labeling and imaging with multiple probes are necessary to determine the presence or absence of specific transcripts in distinct neuronal and glia populations and characterize complex subpopulations Imaging specimens with a large number of fluorescent labels and high autofluorescence often leads to bleed-through or cross-talk of their emission spectra. These artifacts generally complicate the interpretation of experimental findings. To this end, it is necessary to have well-separated fluorescence labels and robust spectral unmixing methods in fluorescence imaging of brain tissue. Given these datasets are becoming increasingly large and necessary to precisely quantify (17), decoupling data collection from linear unmixing and integrating unmixing pipelines with segmentation and quantification tools would increase throughput in acquisition and analysis workflows.

In addition to linear unmixing, many other computational methods are used by biologists for spectral unmixing. Non-negative matrix factorization (NMF) is a feature extraction method that has been successfully applied to unmix multispectral images and does not require prior knowledge of spectral signatures (18,19). However, on different runs, NMF produces equally valid yet significantly different solutions (20). Spectral deconvolution (21) requires users to acquire and manually select each fluorophore in regions of interest. In this age of rapidly advancing machine learning techniques, several unsupervised machine learning methods have been applied to unmix spectral data. Particularly in the field of remote sensing, studies have used clustering methods to separate geological components, such as sand, water, vegetation, etc., in hyperspectral images (22–24). In a recent biological study, McRae et al. used *k*-means clustering to unmix individual pixels in two-photon laser scanning microscopy images (9). Although this method does not require prior information about reference spectra, the performance is sensitive to data preprocessing and robust initialization of cluster centers. Furthermore, this method fails to separate fluorophores that are substantially overlapping. Finally, plug-in options exist in FIJI/ImageJ (25,26) to unmix multispectral images (9,27), but these are limited in terms of handling autofluorescence and biologically similar structures, such as puncta belonging to different gene transcripts.

Linear unmixing assumes a Linear Mixture Model (LMM), where contributions from each endmember sum linearly (3). Although LMM performs very well in most scenarios and reduces computational complexity, the assumption of linearity may be inappropriate in non-linear effects such as quenching and photobleaching. Non-linear unmixing has also been explored in the field of remote sensing (28,29), but both groups of methods fail to account for spectral variability. Spectral signatures deviate from the reference spectra for various reasons, including interaction with surrounding pixels, scattering in complex tissues, autofluorescence, etc. Several methods have been developed to address the shortcomings of LMM in remote sensing data (30,31), but these strategies have not yet been applied to fluorescence microscopy data.

We developed an intuitive and automated pipeline to address the need for improved flexibility, accuracy, and throughput of fluorescence imaging spectral unmixing of brain tissues. Notably, we automate the process of endmember selection using Vertex Component Analysis (VCA) (32). We then use the estimated endmembers to unmix fluorescence images with various unmixing methods derived from remote sensing and hyperspectral imaging. We validate the proposed methods’ accuracy by comparing the unmixing results obtained using ZEN Black software from Zeiss. We also provide scripts to run unmixing in parallel on several images using a High-Performance Cluster (HPC). Finally, we integrate our unmixing pipeline into the computational tool *dotdotdot* (17) to provide a one-stop solution for analysis and quantification of smFISH fluorescence imaging data from mouse and postmortem human brain specimens.

## Materials and methods

### Sample preparations

#### Animals

Wild-type mice were purchased from Jackson laboratories (Bar Harbor, ME, C57BL6/J; stock #000664). All mice were housed in a temperature-controlled environment with a 12:12 light/dark cycle and *ad libitum* access to standard laboratory chow and water. All experimental animal procedures were approved by the JHU Institutional Animal Care and Use Committee.

#### RNAscope single molecule fluorescent *in situ* hybridization (smFISH) for cultured neurons

Mouse cortical neurons were cultured on a 96-well ibidi optical bottom culture plate as previously described (33,34). *In situ* hybridization assays were performed with RNAscope technology using the RNAscope Fluorescent Multiplex kit V2 and 4-plex Ancillary Kit (catalog numbers 323100, 323120 ACD, Hayward, CA) as above, with the exception of the protease step, where cultured cells were treated with a 1:15 dilution of protease III for 30 minutes. Cells were incubated with probes for Arc, Bdnf exon 1, Bdnf exon 4, and Fos (catalog number 316911, 457321-c2, 482981-c3 and 316921-c4, ACD, Hayward, CA) and stored overnight in a 4X saline sodium citrate (SSC) buffer. After amplification, probes were fluorescently labeled with Opal dyes (PerkinElmer; Opal 520 was diluted 1:500 and assigned to *Fos*, Opal 570 was diluted 1:500 and assigned to *Bdnf* exon 1, Opal 620 was diluted 1:500 and assigned to *Bdnf* exon4 and Opal 690 was diluted 1:500 and assigned to *Arc*) and stained with DAPI (4′,6-diamidino-2-phenylindole) to label nuclei, then stored in phosphate buffered saline (PBS) at 4°C.

#### RNAscope smFISH for mouse brain tissue

Mouse brain was extracted and rapidly frozen in 2-methylbutane (ThermoFisher), and stored at −80°C until slicing. Sixteen µm coronal sections were prepared using a Leica CM 1520 Cryostat (Leica Biosystems, Buffalo Grove, IL) and mounted onto glass slides (VWR, SuperFrost Plus). RNAscope was performed using the Fluorescent Multiplex Kit V2 (Cat # 323100, 323120 ACD, Hayward, California) according to manufacturer’s instructions. Briefly, brain sections were fixed in 10% buffered formalin solution (Cat # HT501128 Sigma-Aldrich, St. Louis, Missouri) for 30 min at RT, washed with 1 X PBS, and dehydrated with serial 50%, 70% and 100% ethanol washes. Following pretreatment with hydrogen peroxide, slices were treated with protease IV solution for 20 min and incubated with specific probes targeting *Gal*, *Th, Bdnf* exon 9, and *Npy* (Cat # 400961, 317621-C2, 482981-C3, 313321-C4, ACD, Hayward, California) at 40°C for 2 h in a HybEZ oven (ACD, Hayward, California). Slices were kept in 4X SSC overnight at 4°C and, on the following day, were incubated at 40°C with a series of fluorescent Opal Dyes (Perkin Elmer; Opal690 diluted at 1:500 and assigned to *Npy*; Opal570 diluted at 1:500 and assigned to *Th*; Opal620 diluted at 1:500 and assigned to *Bdnf*; Opal520 diluted at 1:500 and assigned to *Gal*). DAPI was used to label nuclei and slides were coverslipped with FluoroGold (SouthernBiotech).

#### Immunofluorescence staining in post-mortem human Alzheimer’s Disease (AD) brain

Post-mortem human brain tissue was obtained by autopsy from the Offices of the Chief Medical Examiner of Maryland, all with informed consent from the legal next of kin collected under State of Maryland Department of Health and Mental Hygiene Protocol 12-24. Clinical characterization, diagnoses, and macro- and microscopic neuropathological examinations were performed on all samples using a standardized paradigm. Details of tissue acquisition, handling, processing, dissection, clinical characterization, diagnoses, neuropathological examinations, and quality control measures have been described previously (35). Alzheimer’s disease diagnosis comprise standard neuropathology ratings of Braak staging schema (36) evaluating neurofibrillary tangle burden, and the CERAD scoring measure of senile plaque burden (37). An Alzheimer’s likelihood diagnosis was then performed based on the published consensus recommendations for postmortem diagnosis of Alzheimer’s disease [2] as with prior publications (38,39).

Fresh frozen inferior temporal cortex from a donor with clinically confirmed Alzheimer’s disease (AD) was sectioned at 10μm and stored at −80°C. Immunofluorescence staining was performed following a demonstrated protocol provided by 10x Genomics available online (CG000312, 10X Genomics, Pleasanton, California). Briefly, slides were thawed for 1 minute at 37°C and fixed with pre-chilled methanol (Cat #34860, Sigma-Aldrich, St. Louis, Missouri) for 30 minutes at −20°C. Sections were blocked with Human TruStain FcX (Cat #422301, Biolegend, San Diego, California) and 2% BSA (Cat #130-091-376, Miltenyi Biotec, Auburn, California) diluted in Blocking Buffer for 5 minutes at room temperature (RT). Primary antibodies were added in Antibody Diluent (3X SSC, 2% BSA and 0.1% TritonX-100 in nuclease free water) and incubated for 30 minutes at RT. All primary antibodies were diluted from each stock solution at a concentration of 1:100: mouse anti-beta-amyloid (Cat #803001, Biolegend, San Diego, California), rabbit anti-pTau Ser202/Thr205 (Cat # SMC-601, StressMarq Biosciences, Cadboro Bay, Victory, Canada), and chicken anti-MAP2 (Cat #ab92434, Abcam, Cambridge, Massachusetts). The slides were subjected to subsequent 5 washes, each of which takes 30 seconds with Wash Buffer (3X SSC, 2% BSA and 0.1% TritonX-100 in nuclease free water). The tissue sections were then incubated with corresponding fluorescently labeled secondary antibodies diluted from each stock solution at a concentration of 1:500 for 30 minutes at RT. All secondary antibodies were purchased from Thermo Fisher Scientific (Waltham, Massachusetts): goat anti-mouse IgG (H+L) conjugated to Alexa Fluor 488 (Cat #A-11001), donkey anti-rabbit IgG (H+L) conjugated to Alexa Fluor 555 (Cat #A-31572), and goat anti-chicken IgY (H+L) conjugated to Alexa Fluor 633. DAPI was added to visualize the nuclei. After 5 washes with Wash Buffer, which takes 30 seconds for every round, and subsequent 20 quick immersions in 3X SSC (Millipore-Sigma, S6639L, St. Louis, Missouri), slides were coverslipped in 85% glycerol and stored at 4°C.

#### RNAscope smFISH in postmortem human dorsolateral prefrontal cortex (DLPFC)

Two blocks of fresh frozen dorsolateral prefrontal cortex (DLPFC) from neurotypical control individuals ages 24 and 17 were sectioned at 10μm and stored at −80°C as previously described (17). RNA integrity numbers (RINS) were 8.4 and 8.8, respectively. For post-mortem human studies, *in situ* hybridization assays were performed with RNAscope technology utilizing the RNAscope Fluorescent Multiplex Kit V2 and 4-plex Ancillary Kit (Cat # 323100, 323120 ACD, Hayward, California) according to manufacturer’s instructions. Briefly, tissue sections were fixed with a 10% neutral buffered formalin solution (Cat # HT501128 Sigma-Aldrich, St. Louis, Missouri) for 30 minutes at RT, series dehydrated in ethanol, pretreated with hydrogen peroxide for 10 minutes at RT, and treated with protease IV for 20 minutes. Sections were incubated with probes for *SNAP25*, *SLC17A7*, *GAD1*, and *MBP* (Cat #518851, 415611-C2, 573061-C3, 573051-C4, ACD, Hayward, California) and stored overnight in a 4x SSC buffer. Probes were fluorescently labeled with Opal Dyes (Perkin Elmer, Waltham, MA; Opal690 diluted at 1:1000 and assigned to *SNAP25*; Opal570 diluted at 1:1500 and assigned to *SLC17A7*; Opal620 diluted at 1:500 and assigned to *GAD1*; Opal520 diluted at 1:1500 and assigned to *MBP*) and stained with DAPI to label the nucleus (40).

Two subjects were imaged. For each subject, two cortical strips were tile imaged at 20× to capture layers I to VI. Layer II/II and layer VI were identified by measuring 20–30% and 80–90% of the cortical layer thickness, respectively. This strategy reliability delineated layer II/III and VI across 10 individuals and cortical strips with varying absolute thicknesses. After demarcation of cortical layers, the positions feature in Zen software was used to randomly select six fields per layer per strip (n = 12 layer II/III and n = 12 layer VI in two different cortical strips per subject) for high magnification imaging at 63×.

### Fluorescent Imaging

Lambda stacks were acquired in *z*-series using a Zeiss LSM780 confocal microscope equipped with 20x × 1.4 NA and 63x × 1.4NA objectives, a GaAsP spectral detector, and 405, 488, 555, and 647 lasers. All lambda stacks were acquired with the same imaging settings and laser power intensities. Following image acquisition, lambda stacks in *z*-series were linearly unmixed in ZEN software (weighted; no autoscale) using reference emission spectral profiles previously created in ZEN (see below) and saved as Carl Zeiss Image “.czi” files. Raw lambda stacks were unmixed with *SUFI* and compared to ZEN unmixed results. Single-fluorophore positive fingerprints were generated from samples prepared as above.

### Reference spectral profile creation in ZEN software for validation

Reference emission spectral profiles, or ‘fingerprints’, or ‘endmembers’, were created for each Opal dye in ZEN software as previously described (17). Briefly, four single positive slides were generated in mouse tissue using the RNAscope Fluorescent Multiplex Kit V2 and 4-plex Ancillary Kit (Cat # 323100, 323120 ACD, Hayward, California) and a control probe against the housekeeping gene *POLR2A* according to manufacturer’s instructions as described above. Mouse tissue was used in place of human tissue due to lower tissue autofluorescence (i.e. the absence of confounding lipofuscin signals). For every single positive slide, *POLR2A* was labeled with either Opal520, Opal570, Opal620, or Opal690 dye. A single positive slide was generated for DAPI using the same pretreatment conditions, but the omission of probe hybridization steps. To generate a reference emission spectral profile for lipofuscin autofluorescence, a negative control slide was generated in post-mortem DLPFC tissue using a 4-plex negative control probe against four bacterial genes (Cat #321831, ACD, Hayward, CA) in which all Opal dyes were applied, but no probe signal was amplified.

In a similar approach, reference emission spectral profiles were generated for immunofluorescent staining in the post-mortem human AD brain tissue.. For amyloid plaques and tau tangles, each single positive slide was prepared by labeling β-amyloid (Abeta) or phospho-tau (pTau) with appropriate primary and secondary antibodies conjugated with Alexa fluor (AF)488 and AF555, respectively. A lipofuscin fingerprint was created for human AD brain tissue using a negative control slide treated only with fluorescently labeled secondary antibodies in the absence of primary antibodies. For DAPI-stained nuclei and MAP2-positive neurites (labeled with AF633), single positive slides were generated using mouse brain tissue to avoid lipofuscin autofluorescence.

### Spectral unmixing

Spectral unmixing is the process of decomposing composite multichannel images into spectral profiles and abundances of each endmember in each pixel (2) (41) (42):

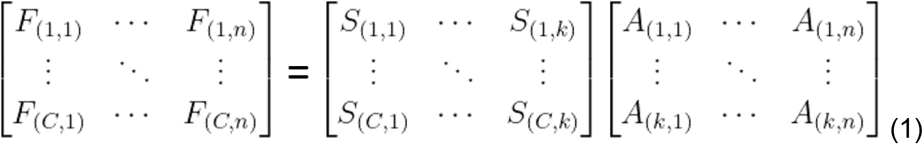

Which can be denoted as F = SA.

In Eq1, F denotes the fluorescence intensities of n pixels recorded in C different spectral channels. S is the spectral signatures of k fluorophores, and A is the abundance of each fluorophore in each pixel. To this end, the unmixing process is usually divided into three different steps: (i) estimation of the number of endmembers, (ii) extraction of endmembers, (iii) estimation of abundance. In fluorescence microscopy, the number of endmembers is known in advance. We discuss the latter two steps below.

#### Automated extraction of spectral signatures

An essential part of the proposed pipeline is the automated extraction of the spectral signatures, or ‘endmembers,’ from the observed multispectral image. To achieve this, we use the Vector Component Analysis (VCA) - an Endmember Extraction Algorithm (32) that can be used to extract fingerprints (i.e. spectral signatures) from multiplex lambda stacks. We approach the extraction of fingerprints in two different ways, (i) Using lambda stacks acquired from the single positive slides from above to extract fingerprints for individual fluorophores. (ii) Using a multiplex lambda stack and extracting fingerprints for all fluorophores in one go. We discuss the pros and cons of each method and provide additional details in the supplementary information.

#### Estimation of abundance

This step involves the estimation of the proportion of different fluorophores in each pixel. Here we implement and compare three different methods derived from remote sensing and adapt them for unmixing in fluorescence microscopy: (i) fully constrained least square unmixing (FCLSU) algorithm (43) tries to minimize the squared error in the linear approximation of multispectral image, imposing the non-negative constraint and the sum-to-one constraint for the abundance calculations. (ii) extended linear mixing model (ELMM) algorithm (30) extends the idea of FCLSU unmixing by taking into account the spectral variability. Particularly, scaling of reference spectra. (iii) generalized extended linear mixing model (GELMM) algorithm (31) extends ELMM to account for complex spectral distortions where different wavelength recordings are affected unevenly.

#### SUFI toolbox

*SUFI* is a MATLAB-based command line toolbox for automated spectral unmixing of fluorescent images. Briefly, the analysis pipeline involves data normalization, automated extraction of spectral signatures using VCA algorithm, and application of spectral unmixing algorithms (Figure 1). Bio-formats toolbox ‘bfmatlab’ is used to read the image data into a MATLAB structure with fields containing gene data, DAPI and lipofuscin. *SUFI* toolbox is publicly available at https://github.com/LieberInstitute/SUFI.

**Figure 1:**
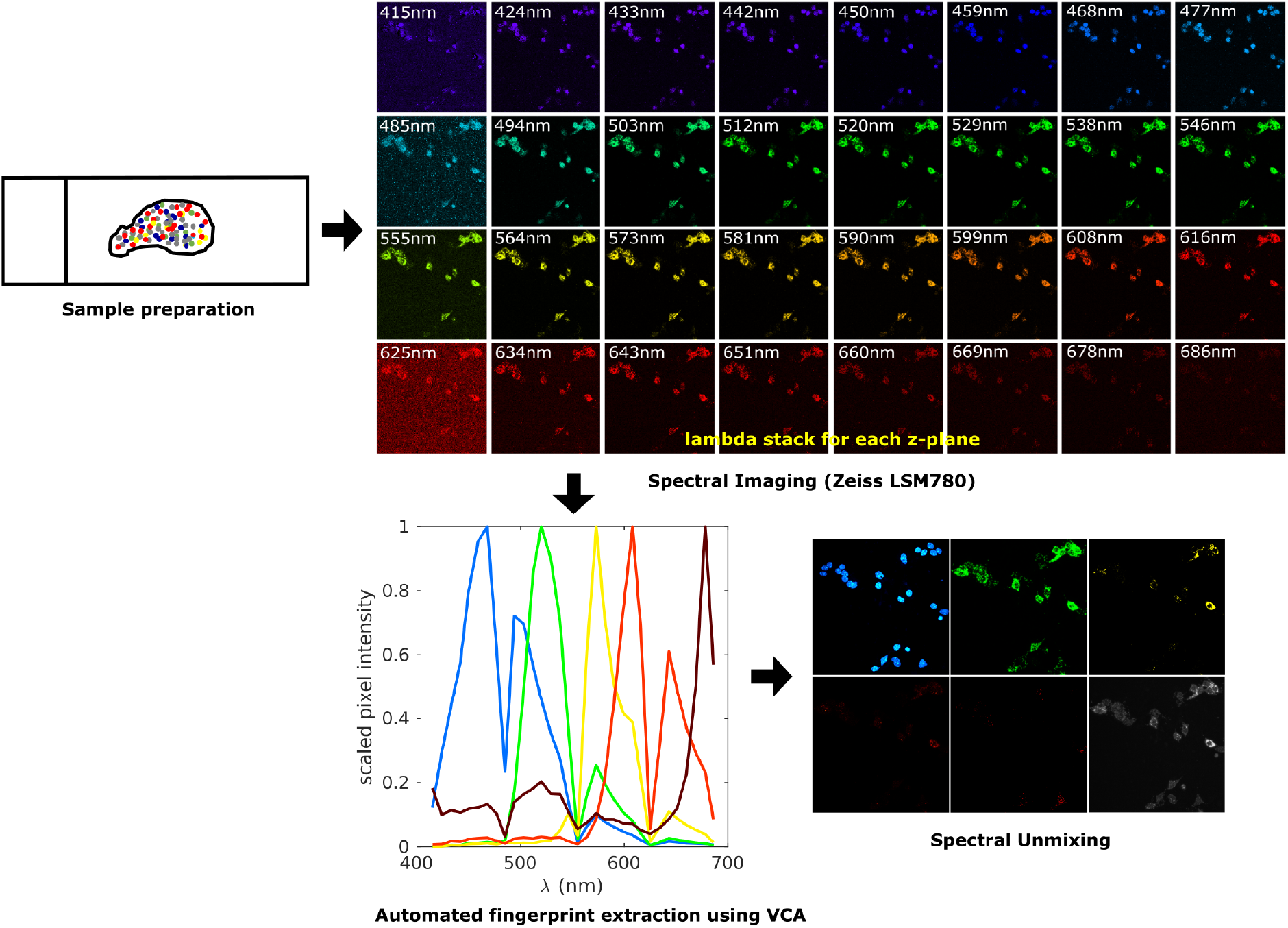
Multispectral imaging and data analysis workflow. Experimental workflow, imaging protocols and data analysis pipelines are similar for mouse and human tissues, but they include optimized conditions for sample preparation (see Methods). Multispectral imaging is performed using a Zeiss LSM780 confocal microscope to acquire lambda stacks. We show a single *z-plane across* the electromagnetic spectrum pseudo colored by wavelength. Spectral signatures are extracted in Matlab using the Vertex Component Analysis. Finally, spectral unmixing is performed using different algorithms to disentangle signals for individual fluorophores.

#### Performance metrics

The Root Mean Squared Error (RMSE) between a true image (y_ref_), i.e. ZEN unmixed image and its estimate (y_est_) i.e. FCLSU (or ELMM or GELMM) unmixed image is defined as,

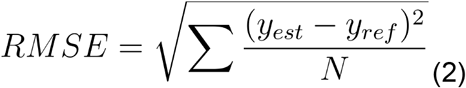

The Structural Similarity Index (SSIM) is based on the computation of luminance, contrast and structure of true image vs. estimated image (44). The range of values are between [0, 1] with a value of SSIM = 1 indicating 100 percent structural similarity.

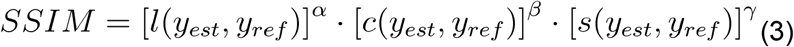

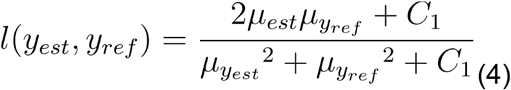

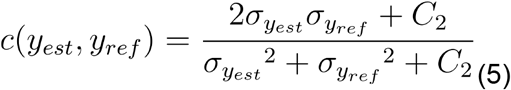

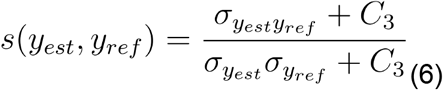

where 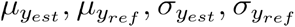, and 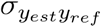 are the local means, standard deviations, and cross-covariance for images *y*_*est*_, *y*_*ref*_. *C*_1_, *C*_2_, *C*_3_ are constants.

The Sørensen-Dice Similarity coefficient (DICE) ranges between [0, 1] where a value of DICE = 1 indicates a 100 percent match of segmentation between two images.

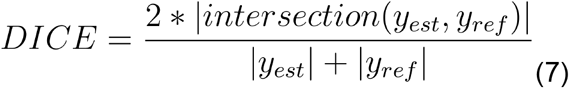

where |*y*_*est*_| represents the cardinal of *y*_*est*_.

### Data segmentation with *dotdotdot*

Using previously published custom MATLAB scripts (17), we automatically segment and quantify nuclei and RNA transcripts using SUFI generated unmixed outputs. Briefly, the *dotdotdot* processing pipeline involves smoothing, thresholding, watershed segmentation, autofluorescence masking, and dot metrics extraction. Specifically, adaptive 3D segmentation is performed on image stacks using the CellSegm MATLAB toolbox, and nuclei are further split using the DAPI channel and 3D watershed function. Single dots are detected using histogram-based thresholding and assigned to nuclei based on their 3D location in the image stack. Lipofuscin signal is used as a mask to remove pixels confounded by autofluorescence.

## Results

In this section, we compare experiment results for four unique biological datasets (cultured mouse neurons, mouse brain tissue, post-mortem human AD brain tissue, and post-mortem human DLFC) using the FCLSU (43), the ELMM (30), and the GELMM (31) unmixing methods. These methods are derived from the field of remote sensing and optimized to our use case. Performances were evaluated in comparison to linearly unmixed images from ZEN Black software (ZEN) using Root Means Squared Error (RMSE), Structural Similarity Index (SSIM), and Sørensen-Dice similarity coefficient (DICE). Runtime values provided are for unmixing on an individual multiplex lambda stack.

### Automated extraction of fingerprints using Vertex Component Analysis

Given the widespread use of multiplex smFISH in human tissues, we first introduce the automated extraction of spectral signatures (i.e., fingerprints) using RNAscope smFISH in postmortem human DLPFC (Figure 2). Single positive images of DAPI, lipofuscin autofluorescence, and RNA transcripts labeled with Opal520, Opal570, Opal620, Opal690 dyes are run through the extraction pipeline. For each single positive lambda stack, two fingerprints are extracted—one representing the fluorophore’s spectral signature and the second representing the background noise’s spectral signature. In order to validate these VCA extracted spectral fingerprints, we manually selected pure pixels from every single positive image and extracted spectral signatures using ZEN software. Plotting VCA-extracted fingerprints against manually extracted ones demonstrates that VCA was able to extract each fingerprint robustly (Figure 2). RMSE values between these curves are calculated and presented within each subplot and further show that the fingerprints extracted through VCA are similar to manually extracted fingerprints.

**Figure 2:**
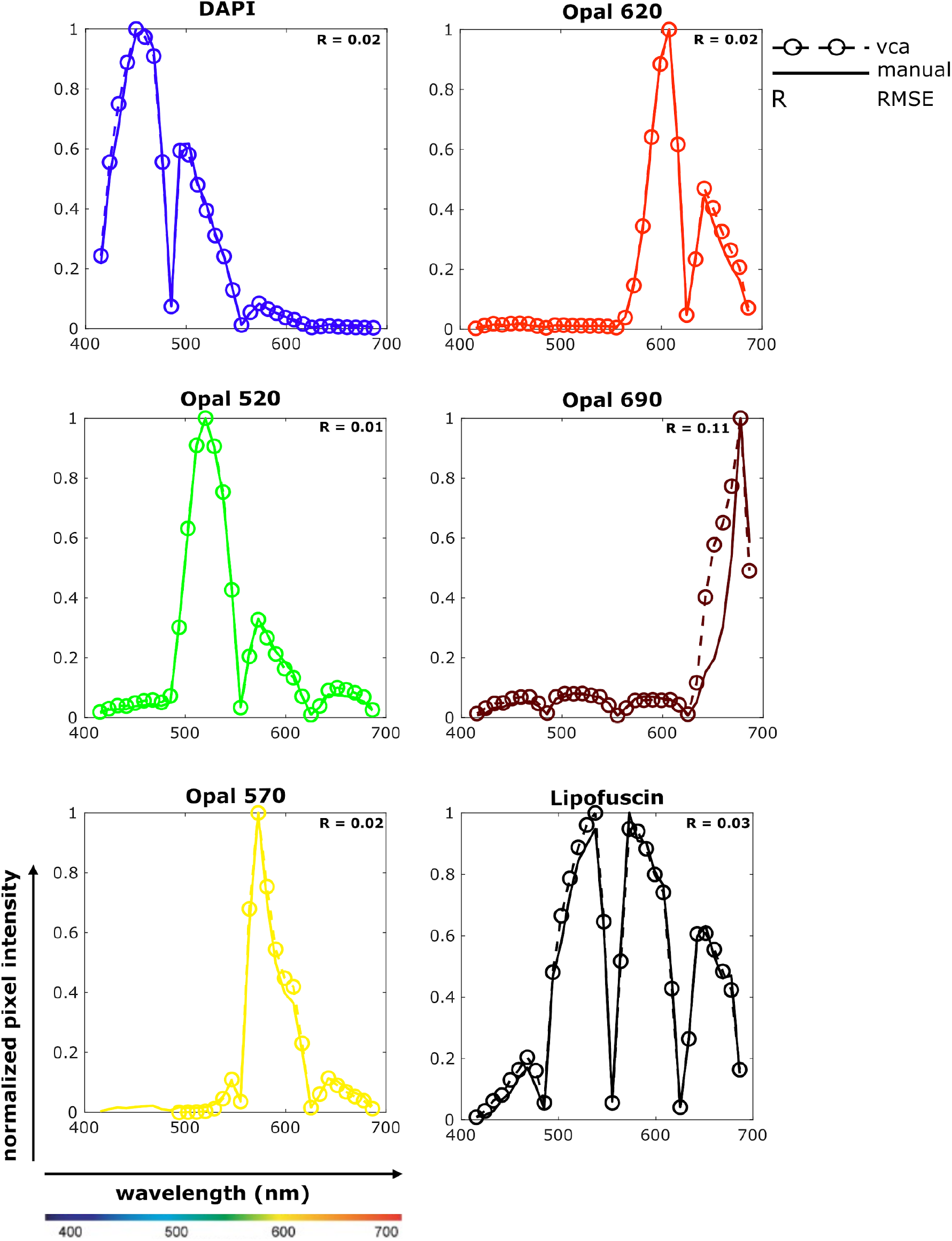
Extracted spectral signatures (fingerprints) from postmortem human brain data. Spectral signatures are extracted from single positive slides using the Vector Component Analysis (VCA). In each subplot, normalized pixel intensity is plotted against wavelength. (i) solid lines represent the fingerprints extracted manually using ZEN Black software. (ii) dotted lines with bubbles represent fingerprints extracted automatically using VCA. The color corresponds to peak wavelength for DAPI and Opal dyes. Lipofuscin is pseudo-colored to black. Root mean squared error (RMSE) between the two lines is calculated for each set of fingerprints

Although it is common to generate single positive samples for spectral imaging experiments, limited reagents or specimen availability may prohibit the generation of single positive slides. To address this, we tried extracting all fingerprints using the multiplex lambda stack instead of lambda stacks acquired from single positive slides. We use the lambda stack from Figure 2 to extract seven pure spectral signatures (i.e., DAPI, Opal520, Opal570, Opal620, Opal690, lipofuscin, and background noise). Plotting and comparing fingerprints extracted from the multiplex lambda stack against single positive extracted fingerprints shows that DAPI and Opal fingerprints are very similar, but the lambda stack approach fails at extracting the lipofuscin fingerprint (Figure S1). Discrepancies for accurately extracting lipofuscin could be due to (i) a smaller fraction of lipofuscin pixels present in the multiplex lambda stack and (ii) the spectral signature of lipofuscin is similar to that of Opal520, thus adding more uncertainty while classifying individual pixels. In summary, we show that VCA can be used as an alternative approach to manual fingerprint generation as long as the fluorophores are spectrally distinct.

### Spectral unmixing of RNAscope multiplexsmFISH for cultured neurons

Having demonstrated successful automated extraction of fingerprints, we first applied reference profiles for unmixing of cultured mouse neuron smFISH data, which lacks strong autofluorescence often observed in tissue slices. These samples contained 5 fluorophores: nuclei stained with DAPI (blue), *Fos* transcripts labeled with Opal520 (green), *Bdnf* exon 1 transcripts labeled with Opal570 (yellow), *Bdnf exon 4* transcripts labeled with Opal 620 (red) and, *Arc* transcripts labeled with Opal690 (maroon). Due to a second peak in the DAPI emission spectrum (Figure 2), the Fos signal in green overlaps with spectrally neighboring blue nuclei signal. We see similar overlap in emission spectra for other combinations of Opal dyes, i.e., *Bdnf* exon 1 in yellow overlapping with *Fos* signal in green, and *Arc* signal in maroon overlapping with *Bdnf* exon 4 in red. Despite spectral overlap in reference emission profiles, all three spectral unmixing algorithms (i.e., FCLSU, ELMM, and GELMM) were able to separate individual fluorophore channels from the multiplex lambda stack (Figure 3). For DAPI, we found that each algorithm performed exceptionally well, scoring 95%+ dice similarity coefficient and low RMSE values (Table 1). We observe similar results for Opal520 with a 97% dice similarity coefficient and even lower RMSE scores (Table 1). Due to the long tail of Opal570 emission spectra, there is some overlap between *Bdnf* exon 1 in yellow and *Bdnf* exon 4 in red (Figure 2). Consequently,the performance of all unmixing algorithms’ is reduced for Opal570 and Opal620 fluorophores (Figure 3C, 3D). Finally, both FCLSU and GELMM accurately unmixed Opal690 signals, scoring 88% dice similarity. On average, FCLSU performed the best, scoring 78% dice similarity with a run time of 10 minutes, GELMM scored 73% dice similarity with a run time of 493 minutes, and ELMM scored 71% dice similarity with a run time of 204 minutes. SSIM remained consistent for each channel across different algorithms.

**Figure 3:**
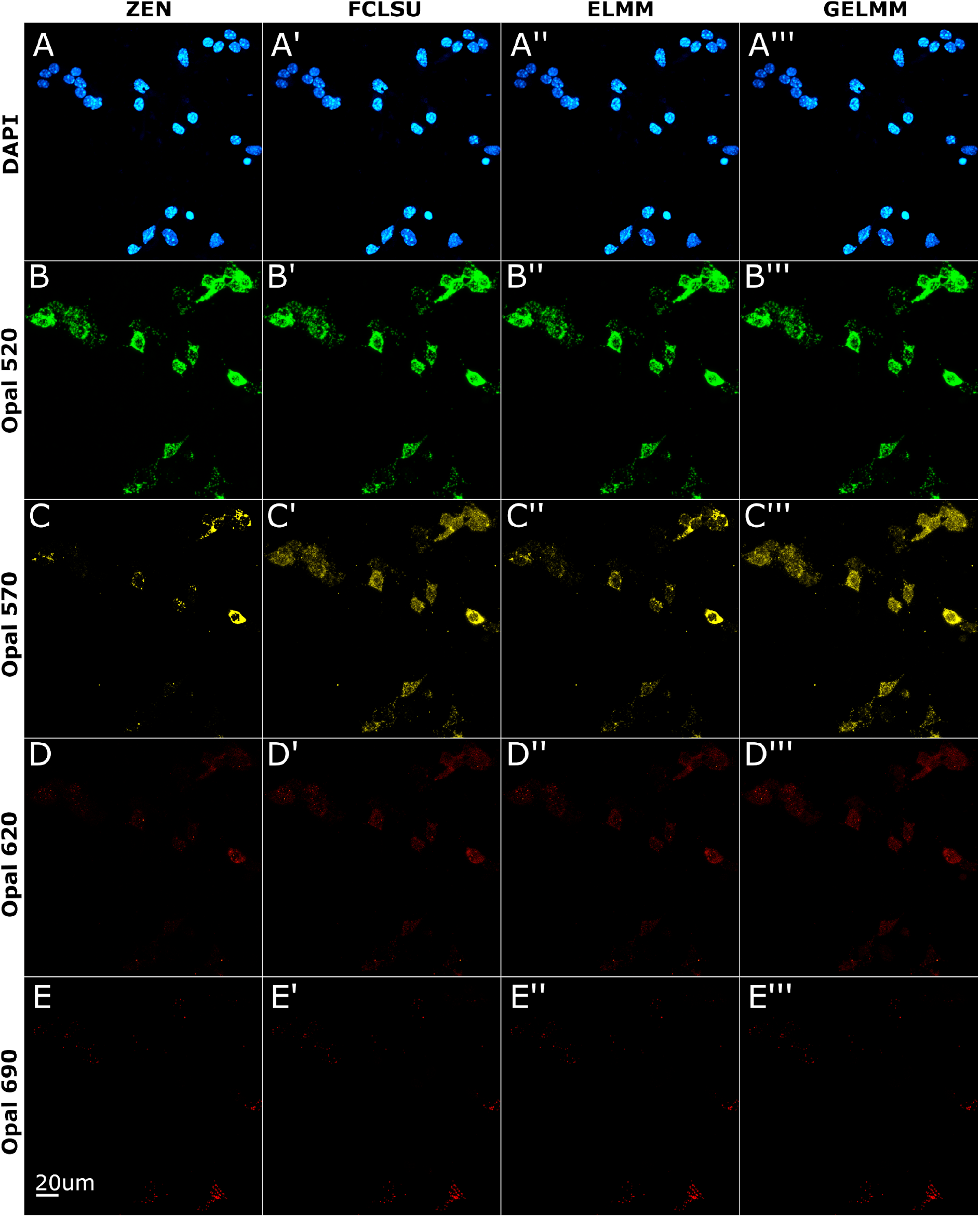
Spectral unmixing of smFISH data from cultured mouse neurons. . Neuronal nuclei stained with DAPI, *Fos* transcripts labeled with Opal520, *Bdnf* exon 1 transcripts labeled with Opal570, *Bdnf* exon 4 transcripts labeled with Opal 620, and *Arc* transcripts labeled with Opal690. Images shown are 2D maximum intensity projections of 3D *z*-stacks. (A - E) Linear unmixing results using ZEN software. (A’ - E’). Spectral unmixing results using FCLSU algorithm. (A’’ - E’’). Spectral unmixing results using ELMM algorithm. (A’’’ - E’’’) Spectral unmixing results using GELMM algorithm.

**Table 1:**
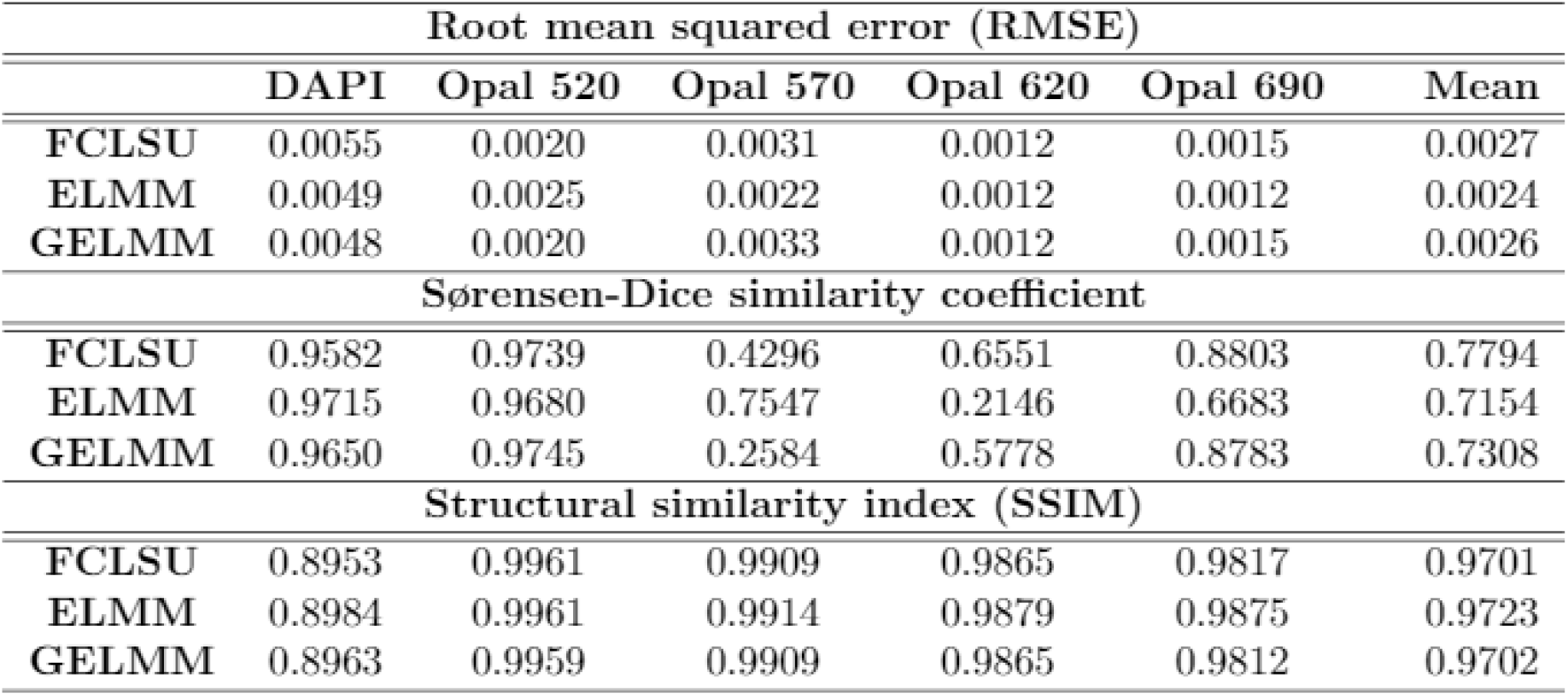
Unmixing results using cultured mouse neuron smFISH data.

| Root mean squared error (RMSE) |  |  |  |  |  |  |
| --- | --- | --- | --- | --- | --- | --- |
|  | DAPI | Opal 520 | Opal 570 | Opal 620 | Opal 690 | Mean |
| **FCLSU** | 0.0055 | 0.0020 | 0.0031 | 0.0012 | 0.0015 | 0.0027 |
| **ELMM** | 0.0049 | 0.0025 | 0.0022 | 0.0012 | 0.0012 | 0.0024 |
| **GELMM** | 0.0048 | 0.0020 | 0.0033 | 0.0012 | 0.0015 | 0.0026 |
| Sørensen-Dice similarity coefficient |  |  |  |  |  |  |
| **FCLSU** | 0.9582 | 0.9739 | 0.4296 | 0.6551 | 0.8803 | 0.7794 |
| **ELMM** | 0.9715 | 0.9680 | 0.7547 | 0.2146 | 0.6683 | 0.7154 |
| **GELMM** | 0.9650 | 0.9745 | 0.2584 | 0.5778 | 0.8783 | 0.7308 |
| Structural similarity index (SSIM) |  |  |  |  |  |  |
| **FCLSU** | 0.8953 | 0.9961 | 0.9909 | 0.9865 | 0.9817 | 0.9701 |
| **ELMM** | 0.8984 | 0.9961 | 0.9914 | 0.9879 | 0.9875 | 0.9723 |
| **GELMM** | 0.8963 | 0.9959 | 0.9909 | 0.9865 | 0.9812 | 0.9702 |

### Spectral unmixing of RNAscope multiplex smFISH for mouse brain tissue slices

Next, we wanted to test the unmixing algorithms on a lower magnification dataset with more densely packed cells. To this end, we applied FLCSU, ELMM, and GELMM algorithms on RNAscope smFISH data collected from mouse brain tissue at 10x magnification. These samples contained 5 fluorophores: cell nuclei stained with DAPI, *Gal* transcripts labeled with Opal520, *Th* transcripts labeled with Opal570, *Bdnf* exon 9 transcripts labeled with Opal620, and *Npy* transcripts labeled with Opal690. Although the unmixed images using the three algorithms look qualitatively similar to unmixed ZEN images (Figure 4), the performance parameters differ significantly (Table 2). This is partly due to densely packed cells resulting in more mixed pixels, thereby increasing the ambiguity in labeling individual pixels. For instance, due to the second smaller peak in the DAPI emission spectrum (Figure 2), the *Gal* signal in green spectrally overlaps with the nuclei signal in blue (Figure 4B - 4B’”). This overlap results in DAPI pixels having spectral signatures of variable amplitude and shape. For DAPI, we see ELMM outperform FCLSU with a dice similarity score of 85% and RMSE of 0.0035 (Table 2). While performance is consistent across different algorithms for Opal520 and Opal570 channels, we see a drop in performance for Opal620 and Opal690 channels with dice similarity around 50% for both fluorophores. On average, ELMM performed the best, scoring 67% dice similarity with a run time of 296 minutes, followed by GELMM scoring65% dice similarity with a run time of 613 minutes, and FCLSU scoring 64% dice similarity with a run time of 6 minutes.

**Figure 4:**
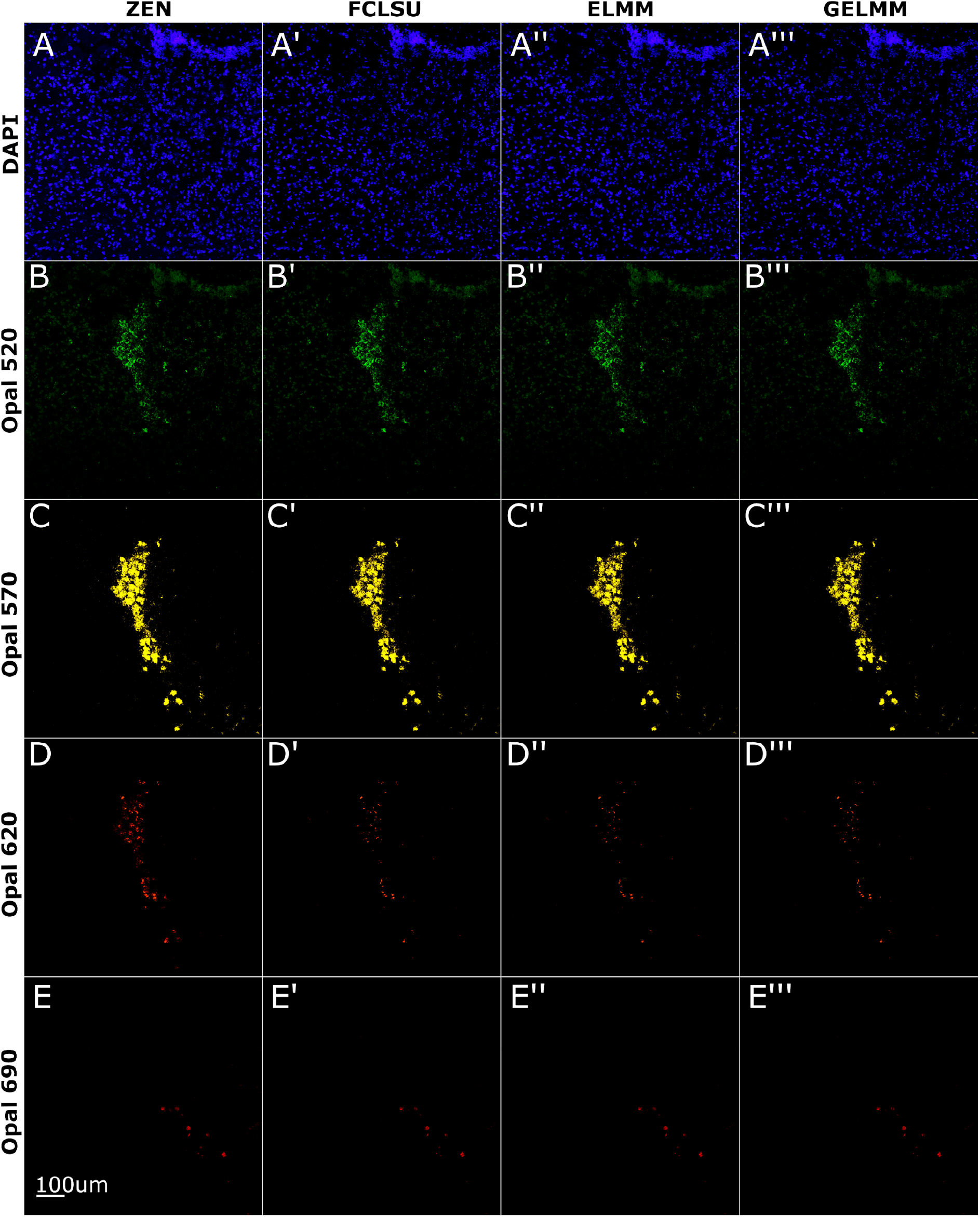
Spectral unmixing of smFISH in mouse brain tissue slices. Nuclei stained with DAPI, *Gal* transcripts labeled with Opal520, *Th* transcripts labeled with Opal570, *Bdnf* exon 9 transcripts labeled with Opal620, and *Npy* transcripts labeled with Opal690. Images shown are 2D maximum intensity projections of 3D *z-*stacks. (A - E) Linear unmixing results using ZEN software. (A’ - E’) Spectral unmixing results using FCLSU algorithm. (A’’ - E’’) Spectral unmixing results using ELMM algorithm. (A’’’ - E’’’) Spectral unmixing results using GELMM algorithm.

**Table 2:** Unmixing results using smFISH data from mouse brain tissue sections.

| Root mean squared error (RMSE) |  |  |  |  |  |  |
| --- | --- | --- | --- | --- | --- | --- |
|  | DAPI | Opal 520 | Opal 570 | Opal 620 | Opal 690 | Mean |
| FCLSU | 0.0052 | 0.0023 | 0.0225 | 0.0079 | 0.0021 | 0.0080 |
| ELMM | 0.0035 | 0.0012 | 0.0248 | 0.0083 | 0.0023 | 0.0080 |
| GELMM | 0.0052 | 0.0021 | 0.0252 | 0.0080 | 0.0021 | 0.0085 |
| Sørensen-Dice similarity coefficient |  |  |  |  |  |  |
| FCLSU | 0.6866 | 0.7574 | 0.8547 | 0.4545 | 0.4752 | 0.6457 |
| ELMM | 0.8559 | 0.8179 | 0.7799 | 0.3918 | 0.5196 | 0.6730 |
| GELMM | 0.7493 | 0.7927 | 0.7795 | 0.4394 | 0.5001 | 0.6522 |
| Structural similarity index (SSIM) |  |  |  |  |  |  |
| FCLSU | 0.8510 | 0.9737 | 0.9682 | 0.9754 | 0.9668 | 0.9470 |
| ELMM | 0.9306 | 0.9963 | 0.9713 | 0.9660 | 0.9580 | 0.9644 |
| GELMM | 0.8466 | 0.9736 | 0.9631 | 0.9748 | 0.9677 | 0.9452 |

### Spectral unmixing of RNAscope multiplex smFISH in postmortem human brain tissue

Autofluorescence is a common, but undesired, signal in fluorescence microscopy that can confound signals from labeled biological targets. Often this background signal has relatively higher intensity and a broader emission spectrum than fluorophores labeling targets (45). Autofluorescence can come from extracellular components or specific cell types and can be more pervasive in certain wavelength bands (46). In human brain tissue, one such autofluorescent material is lipofuscin, a product of lysosomal digestion that accumulates in brain cells due to aging (7) and also in a group of neurodegenerative disorders classified as neuronal ceroid lipofuscinoses, which are characterized by dementia, visual loss, and epilepsy. We next evaluated the ability of the different unmixing algorithms to separate lipofuscin autofluorescence in postmortem human dorsolateral prefrontal cortex (DLPFC) using a previously published dataset (17). This experiment included 5 fluorophores: cell nucleus stained with DAPI, *MBP* transcripts labeled with Opal520, *SLC17A7* transcripts labeled with Opal570, *GAD1* transcripts labeled with Opal620, and *SNAP25* transcripts labeled with Opal690. Like other fluorophores, lipofuscin autofluorescence can be treated as an additional channel and unmixed with its own spectral signature (Figure 2). We see that all unmixing algorithms were able to isolate the autofluorescence signal (Figure 5F’ - 5F’”) and were consistent with high dice similarity scores and low RMSE values (Figure 5, Table 3). Overall, ELMM did best with a dice similarity score of 80% and a runtime of 182 minutes. FCLSU came second with a dice similarity score of 78% and a runtime of 6 minutes. GELMM scored a dice similarity score of 76% and a runtime of 434 minutes.

**Figure 5:**
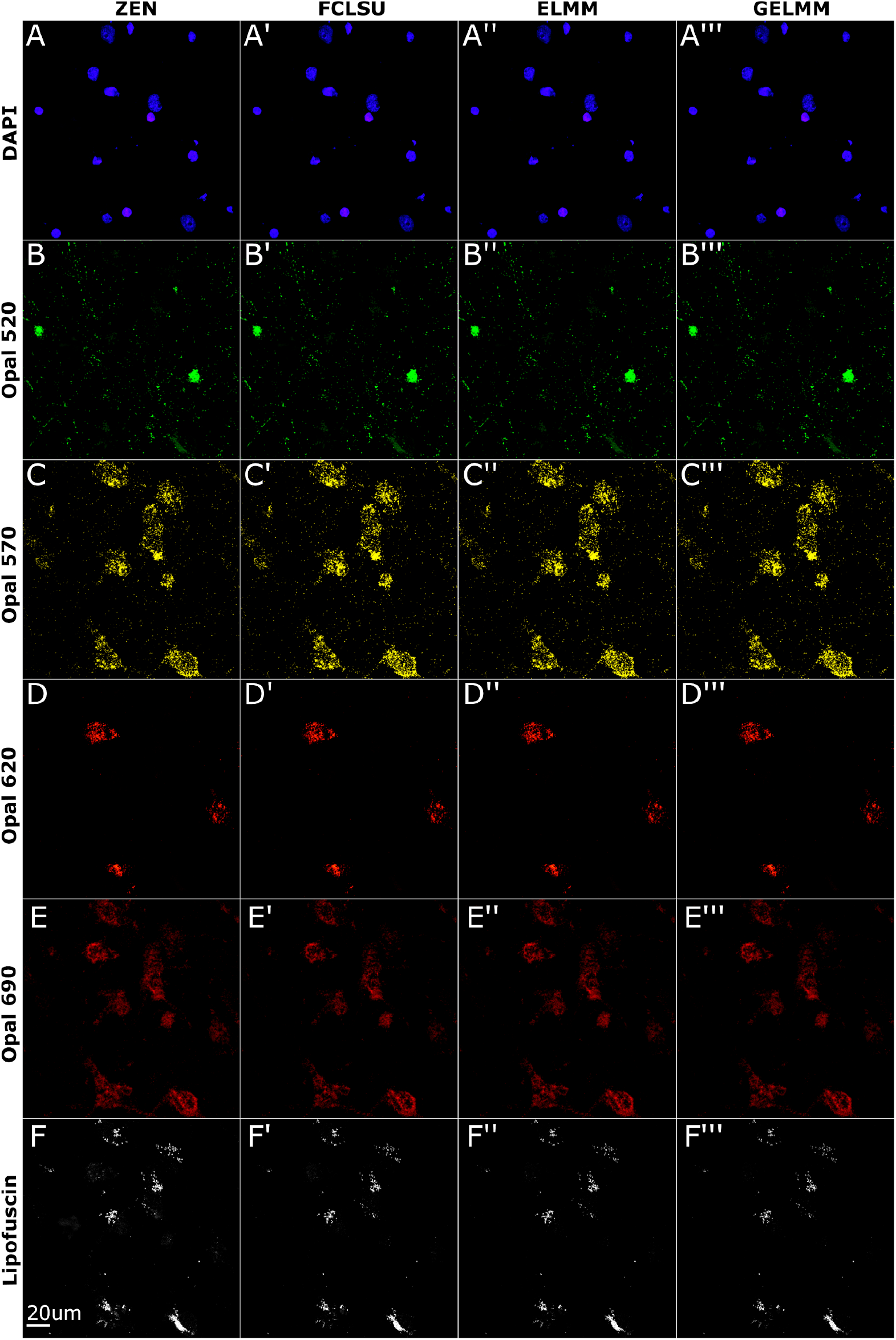
Spectral unmixing of smFISH in postmortem human brain tissue. Cell nucleus is stained with DAPI, *MBP* transcripts labeled with Opal520, *SLC17A7* transcripts labeled with Opal570, *GAD1* transcripts labeled with Opal620, and *SNAP25* transcripts labeled with Opal690. Images shown are 2D maximum intensity projections of 3D *z*-stacks. (A - F) Linear unmixing results using ZEN software. (A’ - F’) Spectral unmixing results using FCLSU algorithm. (A’’ - F’’) Spectral unmixing results using ELMM algorithm. (A’’’ - F’’’) Spectral unmixing results using GELMM algorithm.

**Table 3:**
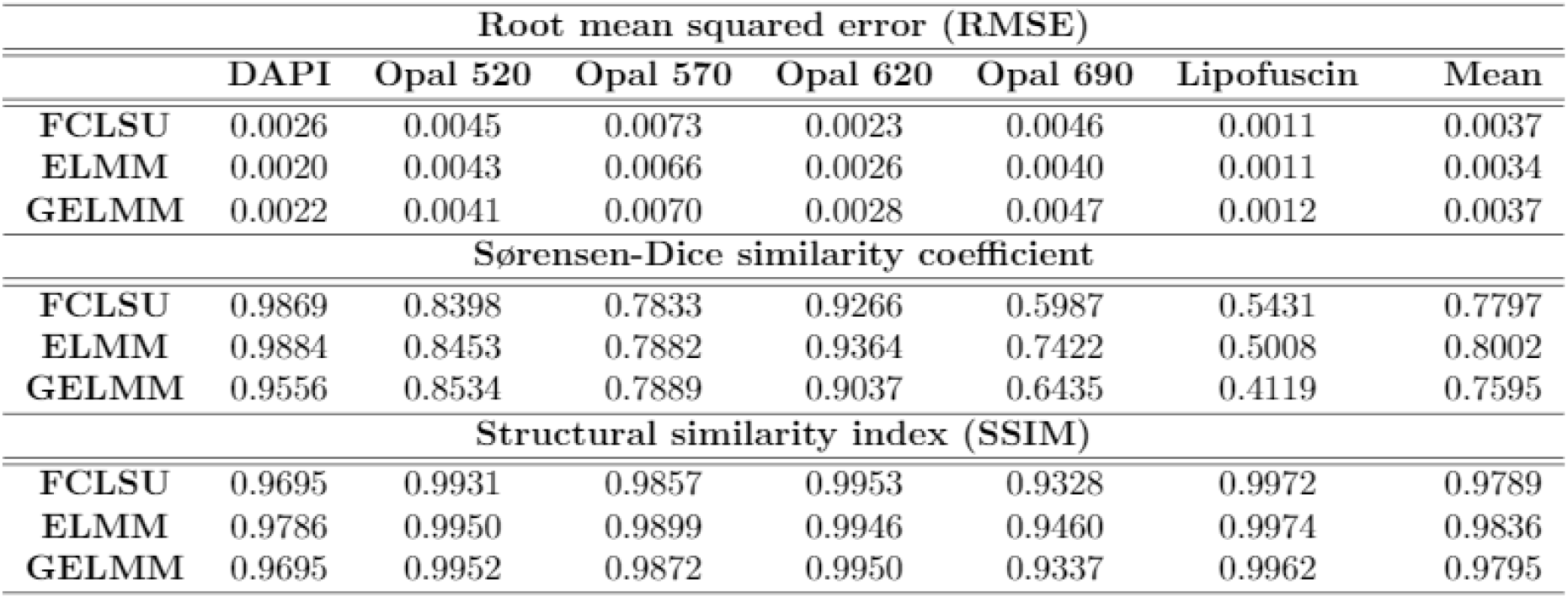
Unmixing results using smFISH data from postmortem human brain tissue sections.

| Root mean squared error (RMSE) |  |  |  |  |  |  |  |
| --- | --- | --- | --- | --- | --- | --- | --- |
|  | DAPI | Opal 520 | Opal 570 | Opal 620 | Opal 690 | Lipofuscin | Mean |
| **FCLSU** | 0.0026 | 0.0045 | 0.0073 | 0.0023 | 0.0046 | 0.0011 | 0.0037 |
| **ELMM** | 0.0020 | 0.0043 | 0.0066 | 0.0026 | 0.0040 | 0.0011 | 0.0034 |
| **GELMM** | 0.0022 | 0.0041 | 0.0070 | 0.0028 | 0.0047 | 0.0012 | 0.0037 |
| Sørensen-Dice similarity coefficient |  |  |  |  |  |  |  |
| **FCLSU** | 0.9869 | 0.8398 | 0.7833 | 0.9266 | 0.5987 | 0.5431 | 0.7797 |
| **ELMM** | 0.9884 | 0.8453 | 0.7882 | 0.9364 | 0.7422 | 0.5008 | 0.8002 |
| **GELMM** | 0.9556 | 0.8534 | 0.7889 | 0.9037 | 0.6435 | 0.4119 | 0.7595 |
| Structural similarity index (SSIM) |  |  |  |  |  |  |  |
| **FCLSU** | 0.9695 | 0.9931 | 0.9857 | 0.9953 | 0.9328 | 0.9972 | 0.9789 |
| **ELMM** | 0.9786 | 0.9950 | 0.9899 | 0.9946 | 0.9460 | 0.9974 | 0.9836 |
| **GELMM** | 0.9695 | 0.9952 | 0.9872 | 0.9950 | 0.9337 | 0.9962 | 0.9795 |

**Table 4:**
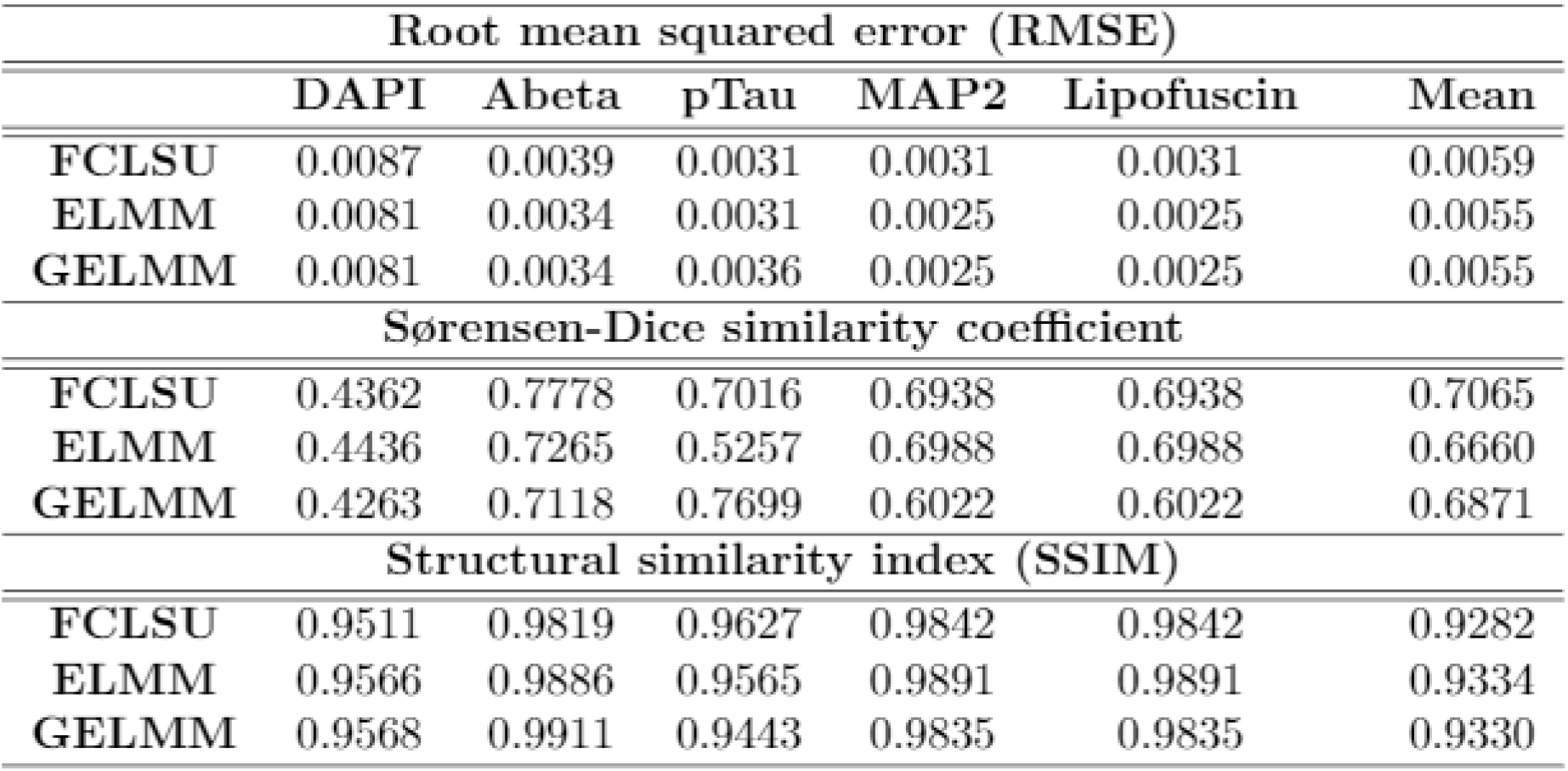
Unmixing results using immunofluorescence data from postmortem human brain tissues sections from AD donors.

| Root mean squared error (RMSE) |  |  |  |  |  |  |
| --- | --- | --- | --- | --- | --- | --- |
|  | DAPI | Abeta | pTau | MAP2 | Lipofuscin | Mean |
| FCLSU | 0.0087 | 0.0039 | 0.0031 | 0.0031 | 0.0031 | 0.0059 |
| ELMM | 0.0081 | 0.0034 | 0.0031 | 0.0025 | 0.0025 | 0.0055 |
| GELMM | 0.0081 | 0.0034 | 0.0036 | 0.0025 | 0.0025 | 0.0055 |
| Sørensen-Dice similarity coefficient |  |  |  |  |  |  |
| FCLSU | 0.4362 | 0.7778 | 0.7016 | 0.6938 | 0.6938 | 0.7065 |
| ELMM | 0.4436 | 0.7265 | 0.5257 | 0.6988 | 0.6988 | 0.6660 |
| GELMM | 0.4263 | 0.7118 | 0.7699 | 0.6022 | 0.6022 | 0.6871 |
| Structural similarity index (SSIM) |  |  |  |  |  |  |
| FCLSU | 0.9511 | 0.9819 | 0.9627 | 0.9842 | 0.9842 | 0.9282 |
| ELMM | 0.9566 | 0.9886 | 0.9565 | 0.9891 | 0.9891 | 0.9334 |
| GELMM | 0.9568 | 0.9911 | 0.9443 | 0.9835 | 0.9835 | 0.9330 |

### Spectral unmixing of immunofluorescence data from postmortem human Alzheimer’s disease brain tissue

Having established that all algorithms performed robustly on different biological samples subjected to smFISH, we lastly wanted to explore the performance of the three algorithms on a different type of fluorescent data acquired using immunofluorescence staining of postmortem human Alzheimer’s disease brain tissue. In this experiment, we also used a different set of fluorophores (Alexa dyes) that were assigned as follows to label nuclei, amyloid beta plaques, neurofibrillary tangles, and neuronal dendrites: the cell nuclei are stained with DAPI, *Abeta* protein is labeled with Alexa fluor (AF)488, *phoso-Tau* protein is labeled with AF555, and *MAP2* protein is labeled with AF633. Spectral signatures for these dyes extracted using VCA are reported in Figure S2. Similar to smFISH data acquired with Opal dyes, we found each algorithm to be robust at unmixing individual fluorophore channels and autofluorescence for immunofluorescence data acquired with Alexa dyes (Figure 6). In terms of performance metrics, pixel-wise RMSE values are consistently under 0.01 (Table 3) andice similarity scores are consistent across different channels and algorithms. Overall, FCLSU performed best with a dice similarity score of 70% and a runtime of 4 minutes. GELMM followed with a dice similarity score of 68% and a runtime of 140 minutes. Finally, ELMM scored a dice similarity score of 66% and a runtime of 41 minutes.

**Figure 6:**
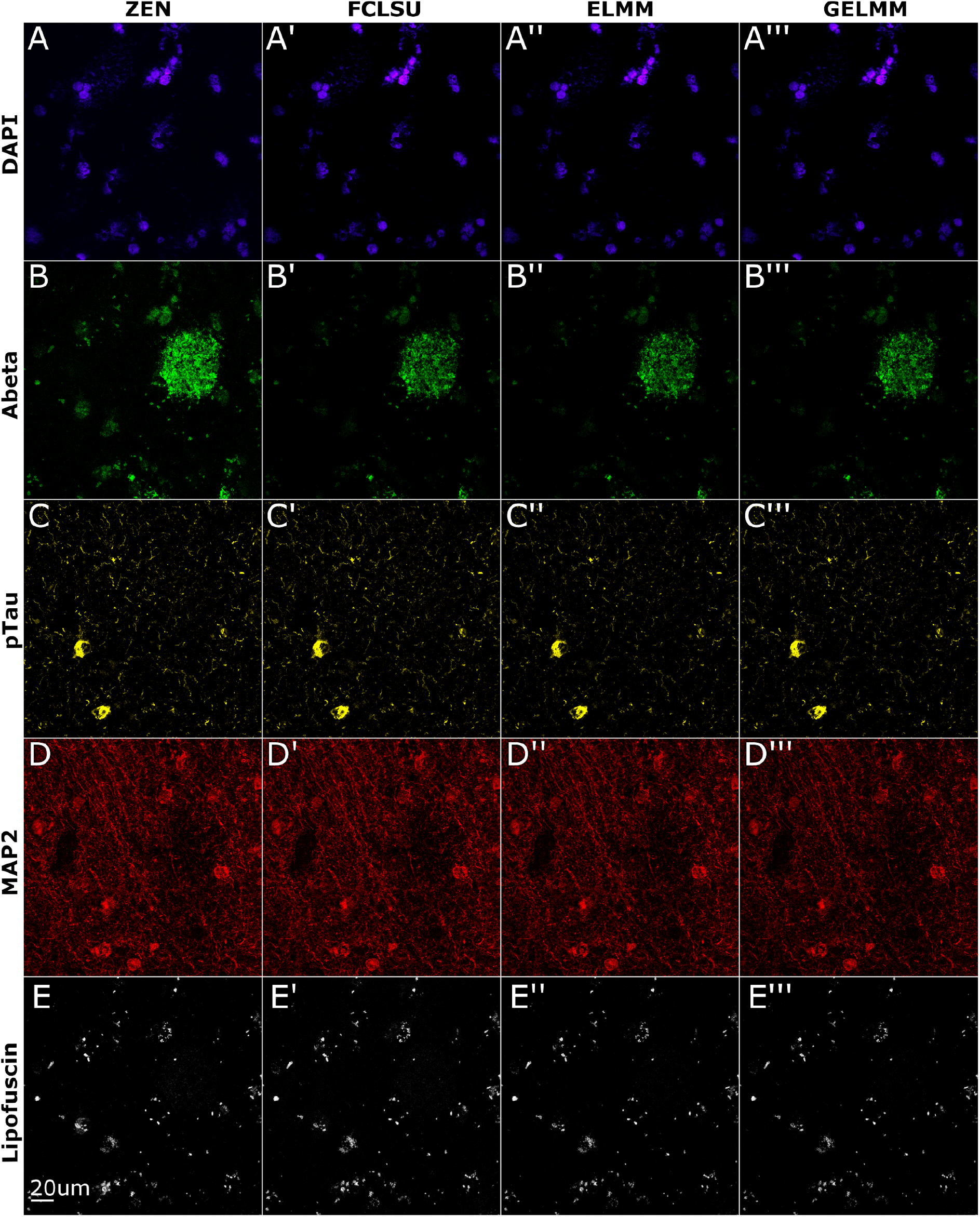
Spectral unmixing of immunofluorescence data from postmortem Alzheimer’s disease brain tissue. The cell nuclei are stained with DAPI, Abeta protein is labelled with AF488, pTau protein is labeled with AF555, MAP2 protein is labeled with AF633. Images shown are 2D maximum intensity projections of 3D *z*-stacks. (A - E) Linear unmixing results using ZEN software. (A’ - E’) Spectral unmixing results using FCLSU algorithm. (A’’ - E’’) Spectral unmixing results using ELMM algorithm. (A’’’ - E’’’) Spectral unmixing results using GELMM algorithm.

### Data segmentation and quantitative analysis using *dotdotdot* framework

Finally, to evaluate the accuracy of other unmixing methods compared to ZEN software, we integrate our automated unmixing pipeline with our previously published *dotdotdot* image segmentation and analysis framework for smFISH data(17). First, we apply FCLSU, ELMM, GELMM, and ZEN-based unmixing on raw smFISH images of postmortem human dorsolateral prefrontal cortex (DLPFC) using the same dataset from Maynard et al. 2020 described in Figure 5 (40). *dotdotdot* automatically segments nuclei and individual transcript channels (Figure 7) and provides several metrics including dot number, dot size, and dot intensity. We quantified the number of segmented objects for each channel in Table 5. For DAPI Opal 520, and Opal 620, we find the number of objects to be equal or consistent across all algorithms. However, we see an increase in the number of objects for Opal 570 when unmixed with FCLSU, ELMM, or GELMM versus ZEN. On the other hand, we see a decrease in the number of segmented objects for Opal 690 and Lipofuscin when unmixed with FCLSU, ELMM, or GELMM versus ZEN. This is potentially due to the proximity of Opal 570 and Opal 690 puncta as seen in Figure 7. Overall, we found the segmentation results of the three different algorithms consistent with those of ZEN-unmixed data (17).

**Figure 7:**
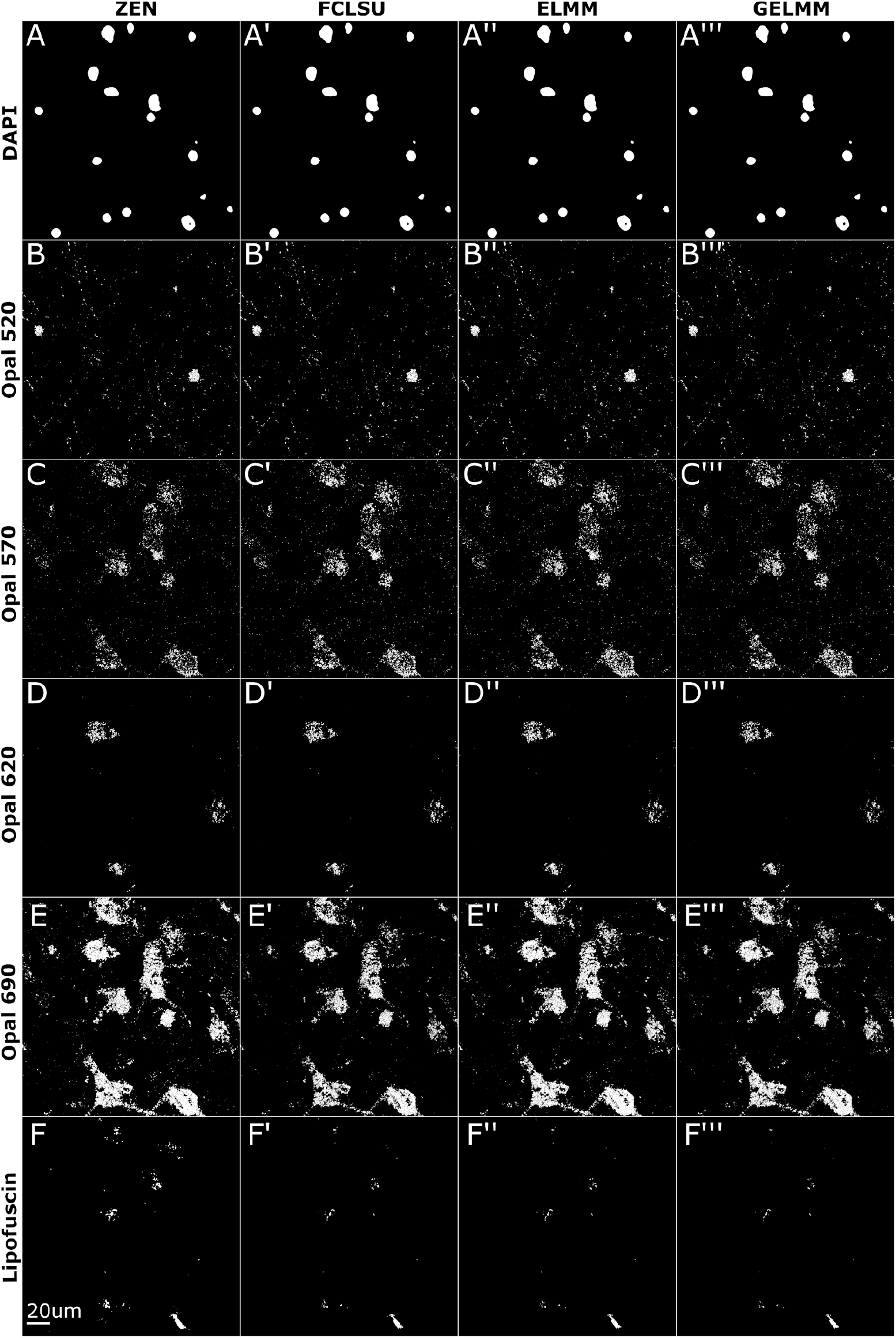
Segmentation of smFISH data in postmortem human brain data using *dotdotdot*.. dotdotdot segmentation of unmixed smFISH images from postmortem human DLPFC. Images shown are 2D maximum intensity projections of 3D segmented z-stacks. (A - F) Segmented results of ZEN unmixed images. (A’ - F’) Segmented results of FCLSU unmixed images. (A’’ - F’’) Segmented results of ELMM unmixed images. (A’’’ - F’’’) Segmented results of GELMM unmixed images.

**Table 5:** Number of segmented objects by *dotdotdot* for human brain tissue data.

|  | DAPI | Opal 520 | Opal 570 | Opal 620 | Opal 690 | Lipofuscin |
| --- | --- | --- | --- | --- | --- | --- |
| ZEN | 17 | 1024 | 2589 | 380 | 8341 | 97 |
| FCLSU | 17 | 1078 | 2907 | 382 | 6564 | 56 |
| ELMM | 17 | 1049 | 2898 | 391 | 7161 | 51 |
| GELMM | 17 | 1028 | 2898 | 380 | 7393 | 43 |

## Discussion

With increasing sophistication in the design and probing of biological targets, there is greater demand for novel imaging technologies that offer enhanced sensitivity, reliable acquisition, and the ability to resolve several targets simultaneously. One such technique, multispectral fluorescence imaging, allows for the observation and analysis of several elements within a sample - each tagged with a different fluorescent dye. Combining multiple fluorescent probes offers a higher level of information from the same sample (47,48) but may also lead to mixed signals (41). Existing spectral unmixing methods solve this problem to some extent, but their accessibility and applicability is limited, especially for smFISH in rodent and brain tissues, and often requires manual intervention. Here, we introduce and systematically evaluate several spectral unmixing algorithms derived from the field of remote sensing that provide a flexible, robust, and automated approach to disentangle multiplex fluorescent images.

We aimed to address several drawbacks to existing spectral unmixing methods for brain tissues that need to be addressed. First, spectral unmixing is often performed using proprietary software that typically accompanies the microscope used for data collection. This leads to several roadblocks regarding cost, throughput, and customization. For instance, it can be challenging to acquire and unmix data on the same computer due to the computational and time requirements, which often necessitates the purchase of additional expensive software licenses. While ZEN software offers a free “lite” version, the full license is required to run unmixing algorithms and each image must be unmixed individually. Also, unmixing algorithms are often not transparent, which leads to a “black-box” in the analysis pipeline that can limit optimization and troubleshooting capabilities. Additionally, spectral unmixing is a computationally-intensive - yet entirely parallelizable - task. The time spent unmixing individual images in between acquisition sessions could instead be directed for additional data collection. For example, in a single imaging session, researchers could spend 3-4hours continuously imaging, which reduces to 2-3 hours of imaging when needing to unmix individual images in real time in a typical RNAscope experiment. The time complexity and inability to unmix in batch mode is incredibly daunting when several hundred images need to be collected and unmixed.

We address these analytical limitations for spectral imaging by providing a MATLAB-based unmixing solution that leverages the implicit parallelizability of unmixing each image independently. By decoupling the data collection and unmixing processes, we aim to improve analysis efficiency. Secondly, unmixing methods based on linear mixing models (LMM), such as linear unmixing, spectral deconvolution, and similarity unmixing, requires the user to choose the individual reference spectrum before unmixing. Manually selecting pure pixels in a single positive or multiplex lambda stack for fingerprint generation is a labor-intensive task that is also prone to human bias since pure pixels are variable between images or even within the same image, which may result in the generation of variable reference profiles. Also, background noise and autofluorescence need their own spectral signatures, which are more challenging to estimate manually. We solve this by automating the extraction of spectral signatures using the vertex component analysis (VCA) algorithm, which alleviates the need to manually select pixels for fingerprint generation. Furthermore, background noise and lipofuscin autofluorescence are separated as individual channels with their own spectral signatures using VCA (Figure 2). Lastly, unmixing methods based on unsupervised machine learning algorithms, like clustering, have been introduced in multispectral fluorescence microscopy (9). Although these methods do not require prior information about the reference spectrum, the performance is sensitive to data conditioning and cluster centers’ initialization, which means that we could end up with different unmixed channels for different cluster center initialization. These approaches also tend to perform worse when substantial spectral overlap exists between fluorophores. On the other hand, the only preprocessing step for our pipeline is data normalization, and our approach handles overlapping fluorophores commonly used for smFISH successfully (Figure 5).

In this age of interdisciplinary research, machine learning methods have bought several seemingly unrelated fields closer together. One such field that shares a similar conceptual framework to spectral unmixing is satellite imaging (22,49,50). Particularly in the field of remote sensing, studies have used spectral unmixing methods to disentangle geological channels, such as sand, water, vegetation, etc, from hyperspectral images captured by satellites (23,24). However, spectral unmixing in fluorescence microscopy has several advantages. First, the number of endmembers (fingerprints or spectral signatures) is known in advance based on the number of fluorophores or dyes utilized by the experimenter. Typically, with remote sensing images, the number of endmembers is not known and often calculated as the first step during the unmixing process. Second, due to the limited spatial resolution of satellite images, a large proportion of individual pixels are a mixture of two or more geological components. However, in fluorescence images, individual pixels specify a single fluorophore unless a colocalized label was designed. Conversely, remote sensing images have the upper hand when it comes to the number of spectral bands. Typically, satellite imagery results in several hundred spectral bands, whereas fluorescence microscopy has 30 - 50 bands. In our imaging setup, data is collected at 32 spectral bands spanning the visual spectrum ~every 8nm. Increasing the number of detectors and therefore increasing the number of spectral bands will improve unmixing performance in fluorescent biological samples.

We acknowledge that there are limitations to the methods introduced in this paper. First, these methods assume a linear mixture model (LMM) to disentangle mixed signals mathematically. Although LMM performs very well in most scenarios and reduces computational complexity, the assumption might be inappropriate when non-linear effects such as quenching and photobleaching are used. Second, our methods require that the number of fluorophores is equal to or lesser than the number of spectral bands. If the number of fluorophores is greater than the number of spectral bands, we end up with an under-determined system that cannot be resolved. Third, any increase in the dimension of images leads to an exponential increase in runtime. We used 1024X1024 images in our experiments and ran all unmixing methods on a 40 member cluster node. On average, FCLSU took 6 minutes, ELMM took 180 minutes, and GELMM took 400 minutes. For most scenarios, FCLSU is a clear choice in terms of performance per time utilized. This is because ELMM and GELMM try to improve on results obtained through FCLSU, which in most cases are sufficient enough. Finally, the automated extraction of fingerprints requires single positive images to determine spectral signatures (Figure 2, S2) accurately. Typically, single positive images are generated to validate fluorophores before spectral imaging. Although we show that fingerprints can be extracted directly from the multiplex lambda stack (FIgure S1), this method fails to extract significantly overlapping fingerprints (i.e., Opal 520 and Lipofuscin). Future studies should evaluate data driven approaches to extract spectral signatures and explore deep learning based methods for spectral unmixing.

In conclusion, we present a robust, flexible, and automated tool for spectral unmixing of fluorescent images acquired in brain tissues. Notably, we automate the process of endmember selection using Vertex Component Analysis (VCA). We then provide several unmixing methods derived from remote sensing to disentangle multiplex lambda stacks. We demonstrate these algorithms using four biologically unique fluorescence imaging datasets. Finally, we provide a MATLAB toolbox that can be readily adopted by the scientific community for their unmixing needs and bundled this pipeline with our previous smFISH segmentation/quantification tool *dotdotdot*.

## Acknowledgments

The authors gratefully acknowledge the contributions of the Offices of the Chief Medical Examiner of Maryland for collaborating in the accession of post-mortem human brain donations that were used in this study. We also thank the members of the Neuropathology Section of the Lieber Institute for Brain Development who made important contributions in the clinical characterization and diagnosis of the donors. We thank Dr. Keri Martinowich for comments on the manuscript.

## Data Availability

All data and software generated or analyzed during this study are available on github https://github.com/LieberInstitute/SUFI

---

**Figure S1:**
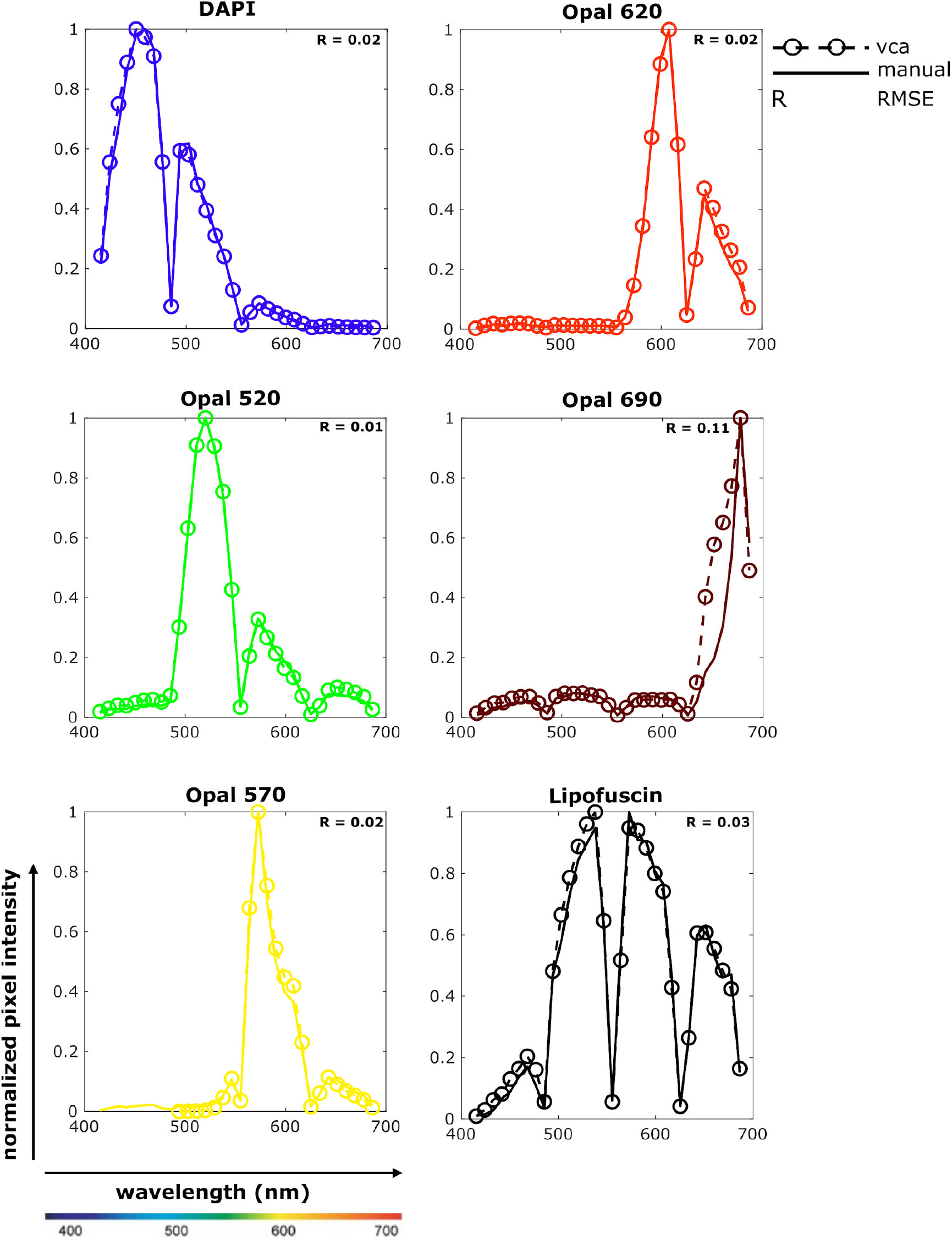
Extracted spectral signatures (fingerprints) from single positive vs. multiplex lambda stacks. Comparison of spectral signatures extracted from single positives vs multiplex lambda stack using the Vector Component Analysis (VCA). In each subplot, normalized pixel intensity is plotted against wavelength. (i) solid lines represent the fingerprints extracted using multiplex lambda stack. (ii) dotted lines with bubbles represent fingerprints extracted from single positives. The color corresponds to peak wavelength for DAPI and Opal dyes. Lipofuscin is pseudo-colored to black. Root mean squared error (RMSE) between the two lines is calculated for each set of fingerprints.

**Figure S2:**
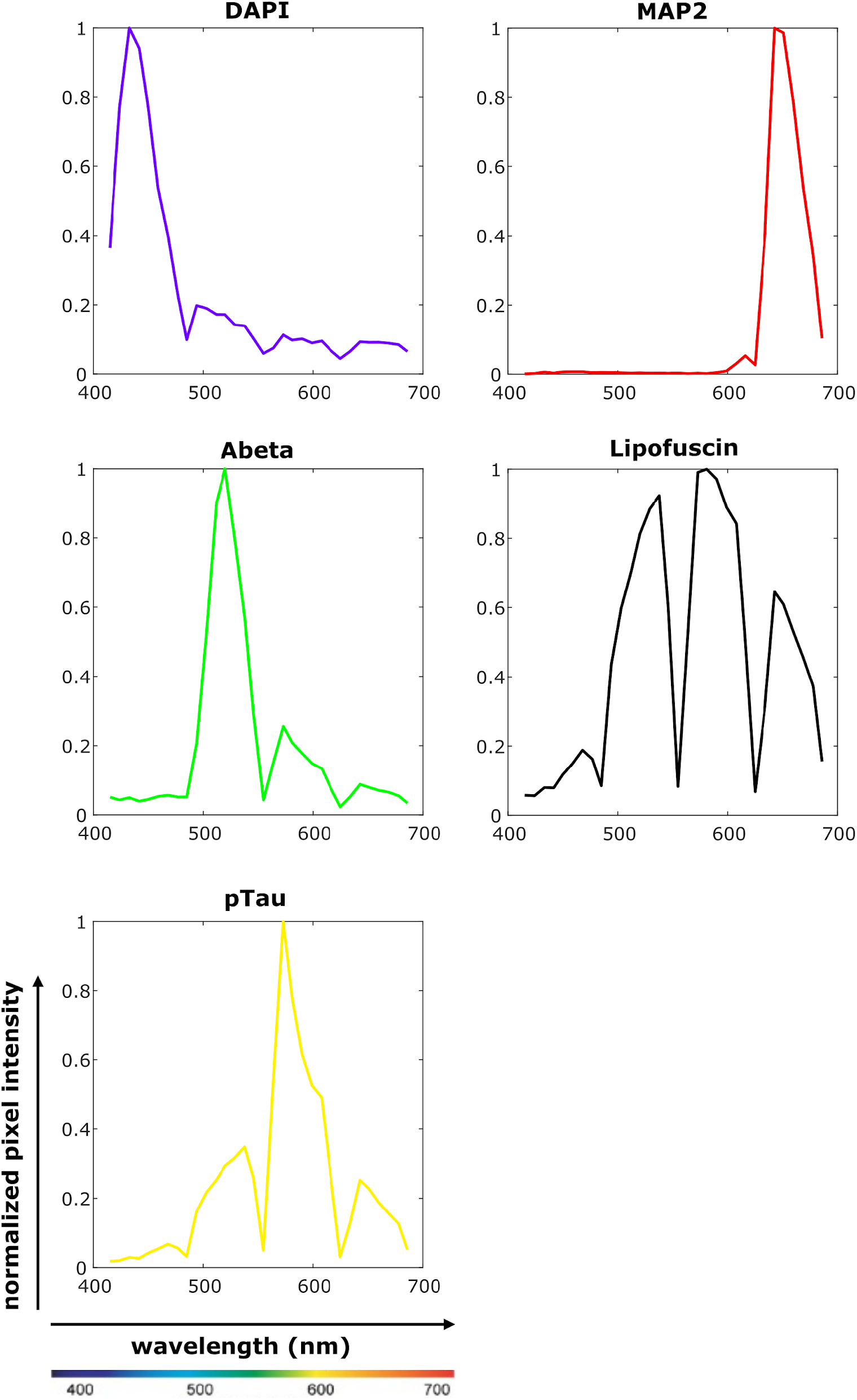
Extracted spectral signatures (fingerprints) of immunofluorescence data from postmortem human brain tissue sections derived from donors with Alzheimer’s disease. Spectral signatures are extracted from single positive lambda stacks using the Vector Component Analysis (VCA). In each subplot, normalized pixel intensity is plotted against wavelength. The color corresponds to peak wavelength for DAPI and Opal dyes. Lipofuscin is pseudo-colored to black.

